# KinFin: Software for taxon-aware analysis of clustered protein sequences

**DOI:** 10.1101/159145

**Authors:** Dominik R. Laetsch, Mark L. Blaxter

**Affiliations:** Institute of Evolutionary Biology, University of Edinburgh, Edinburgh EH9 3JT UK; The James Hutton Institute, Errol Road, Dundee DD2 5DA UK

**Keywords:** bioinformatics, protein orthology, protein family evolution, comparative genomics, filarial nematodes

## Abstract

The field of comparative genomics is concerned with the study of similarities and differences between the information encoded in the genomes of organisms. A common approach is to define gene families by clustering protein sequences based on sequence similarity, and analyse protein cluster presence and absence in different species groups as a guide to biology. Due to the high dimensionality of these data, downstream analysis of protein clusters inferred from large numbers of species, or species with many genes, is non-trivial, and few solutions exist for transparent, reproducible and customisable analyses. We present KinFin, a streamlined software solution capable of integrating data from common file formats and delivering aggregative annotation of protein clusters. KinFin delivers analyses based on systematic taxonomy of the species analysed, or on user-defined groupings of taxa, for example sets based on attributes such as life history traits, organismal phenotypes, or competing phylogenetic hypotheses. Results are reported through graphical and detailed text output files. We illustrate the utility of the KinFin pipeline by addressing questions regarding the biology of filarial nematodes, which include parasites of veterinary and medical importance. We resolve the phylogenetic relationships between the species and explore functional annotation of proteins in clusters in key lineages and between custom taxon sets, identifying gene families of interest. KinFin can easily be integrated into existing comparative genomic workflows and promotes transparent and reproducible analysis of clustered protein data.

## Introduction

Inference of gene homology across taxa is an integral part of comparative genomic analysis. Candidate orthologues, paralogues, and xenologues between taxa are commonly identified through clustering of protein sequence data using tools such as OrthoFinder (Emms and Kelly 2015), OrthoMCL (Li et al. 2003), and others. Exploitation of orthologue definitions across species, the study of gene family evolution, genome evolution, species phylogenetics, and as loci for population genetics and ecological genetics, is demanding. Many research projects aim to identify orthologues of interest that have a specific distribution across species, for example identifying gene families that are synapomorphic for, or that have been specifically lost from, a particular clade. Exploring the effects of assuming different underlying phylogenies on the analysis of the origins of orthologues may assist in discriminating between competing hypotheses. Grouping species by non-phylogenetic classifiers (such as habitat, mating system or life history) may also identify protein families uniquely present/absent or exhibiting differential copy-number.

Several high-quality solutions to orthology analysis have been proposed. OrthoDB is a high- quality curated orthology resource (Zdobnov et al. 2017). The current release (2015) includes 3,600 bacterial and 590 eukaryotic taxa, and is accessed through a responsive web interface or direct download and interrogation. OrthoDB includes rich functional annotation of sequences. While the main database includes only published genomes, and is centrally managed (i.e. users cannot submit datasets for analysis), the OrthoDb software toolkit is available for local installation and deployment. PhylomeDB is a database of defined orthology groups, built with manual curation (Huerta-Cepas et al. 2014), but was last updated in 2014, and is, again, managed centrally and focussed on published genomes. In the ENSEMBL databases, the Compara toolkit is used to parse gene homology sets, and infers orthology and paralogy based on a given species tree (Herrero et al. 2016). Updating of Compara analyses is not trivial, and requires the ENSEMBL web toolkit for display and interrogation. For ongoing research programs, few tools exist for orthology analysis. For bacterial data, several tools for pan-genome analysis have been developed (Vinuesa and Contreras-Moreira 2015; Chaudhari et al. 2016; Xiao et al. 2015) but solutions that cope well with the data richness of eukaryotic species are often tailored to defined taxonomic groups (Song et al. 2015) or expect closely related taxa. EUPAN is a pipeline for pan-genome analysis of closely related eukaryotic genomes developed within the scope of the 3000 Rice Genomes Project (Hu et al. 2017). The approach parts from mapping of raw reads to reference genomes followed by coordinated assembly and lift-over of gene annotations for inferring presence/absence of gene models.

In the absence of toolkits that allow local implementation of clustering analyses, custom taxon grouping and dynamic analysis, we have developed KinFin. KinFin takes a protein clustering output by tools such as OrthoFinder or OrthoMCL, alongside functional annotation data, and user-defined species taxonomies, to derive rich aggregative annotation of orthology groups. KinFin reads from standard file formats produced from commonly-used genome sequencing and annotation projects. KinFin can easily be integrated in comparative genomics projects for the identification of protein clusters of interest in user-defined, taxon-aware contexts.

## Implementation

KinFin is a standalone Python application. A detailed description of the functionality of KinFin can be found at https://kinfin.readme.io/. Required input for KinFin is an orthology clustering (format defined by OrthoMCL/OrthoFinder), a file linking protein sequences to taxa (SequenceIDs defined by OrthoFinder), and a user-defined config file. The config file guides analyses by grouping taxa into user-defined sets under arbitrary attributes. These attributes could include, for instance, standard taxonomy (as embodied in the NCBI Taxonomy “TaxID” identifiers), alternate systematic arrangements of the taxa involved, lifestyle, geographical source or any other aspect of phenotype or other metadata. KinFin dynamically constructs sets based on the config file and computes metrics, statistics and visualisations which allow identification of clusters that are diagnostic for, or expanded/contracted in, each taxon set. Optional input files include proteome FASTA files (to extract length statistics for clusters, taxa and taxon sets), functional annotations of proteins in InterProScan (Jones et al. 2014) format, and a phylogenetic tree topology in Newick format.

### Visualisation of orthologue clustering

In KinFin, global analysis of the clustering of protein sequences can be performed from the point of view of the clusters themselves (their properties and patterns) or of the constituent proteomes. The distribution of cluster size (i.e. the number of proteins contained in a cluster) is an important feature of analyses, and KinFin simplifies the comparison of alternative clusterings (for example ones using different MCL inflation parameters, or with overlapping but distinct taxon inclusion) by generating frequency histograms of cluster size, which can be interrogated for deviations from the expected power law-like distribution. To aid understanding of the distribution, the user can generate a more detailed frequency histogram which considers the number of taxa contributing to each cluster (see Figure 1). The behaviour of individual proteomes can be explored by creating a network representation of the clustering. KinFin can produce a graph file with nodes representing proteomes and edges connecting nodes weighted by the number of times two proteomes co-occur in clusters. Optionally, universal clusters, with members from all the proteomes, can be excluded. The graph can be interrogated using graph analysis and visualisation tools such as Gephi (Bastian et al. 2009) (see Supplementary Figure 1).

**Figure 1.**
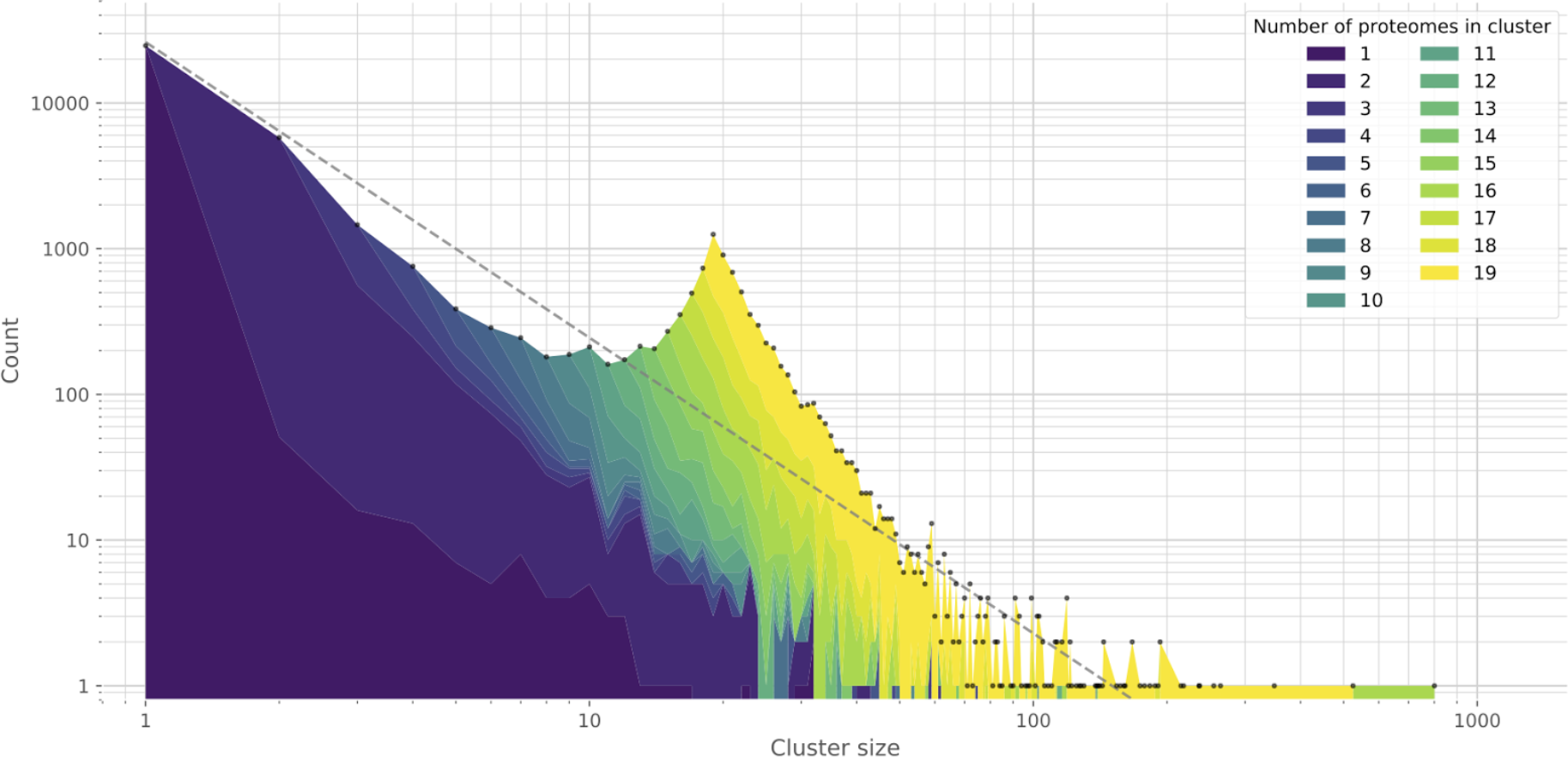
Distribution of cluster sizes (count of proteins per cluster) in clustering. The distribution is coloured based on the number of proteomes present. Total values of counts of each cluster size are indicated with grey dots. A fitted power-law curve (grey) is drawn for reference.

### Analyses based on arbitrary sets of input proteomes

Through the config file, the user can instruct KinFin to analyse the clustering under arbitrary sets. For taxonomy-based analyses, KinFin derives analyses at different taxonomic ranks (by default “phylum”, “order”, and “genus”; can be modified by the user) by parsing the NCBI TaxIDs given for each proteome. Any other classification of the input proteomes can be given, and nested taxonomies specified by use of multiple, ranked attribute types. This allows, for example, the testing of congruence of clustering data with competing phylogenetic hypotheses regarding relationships of the taxa from which the input proteomes were derived.

### Classification of clusters

KinFin builds a series of matrices associating clusters and proteomes, and clusters and arbitrary proteome sets. Each cluster is classified as `absent` or `present` for each proteome or taxon set, and is assigned a cluster type (`singleton`, `specific` – where there are ≥2 members and all come from one taxon set – or `shared`) (see Figure 2).

**Figure 2.**
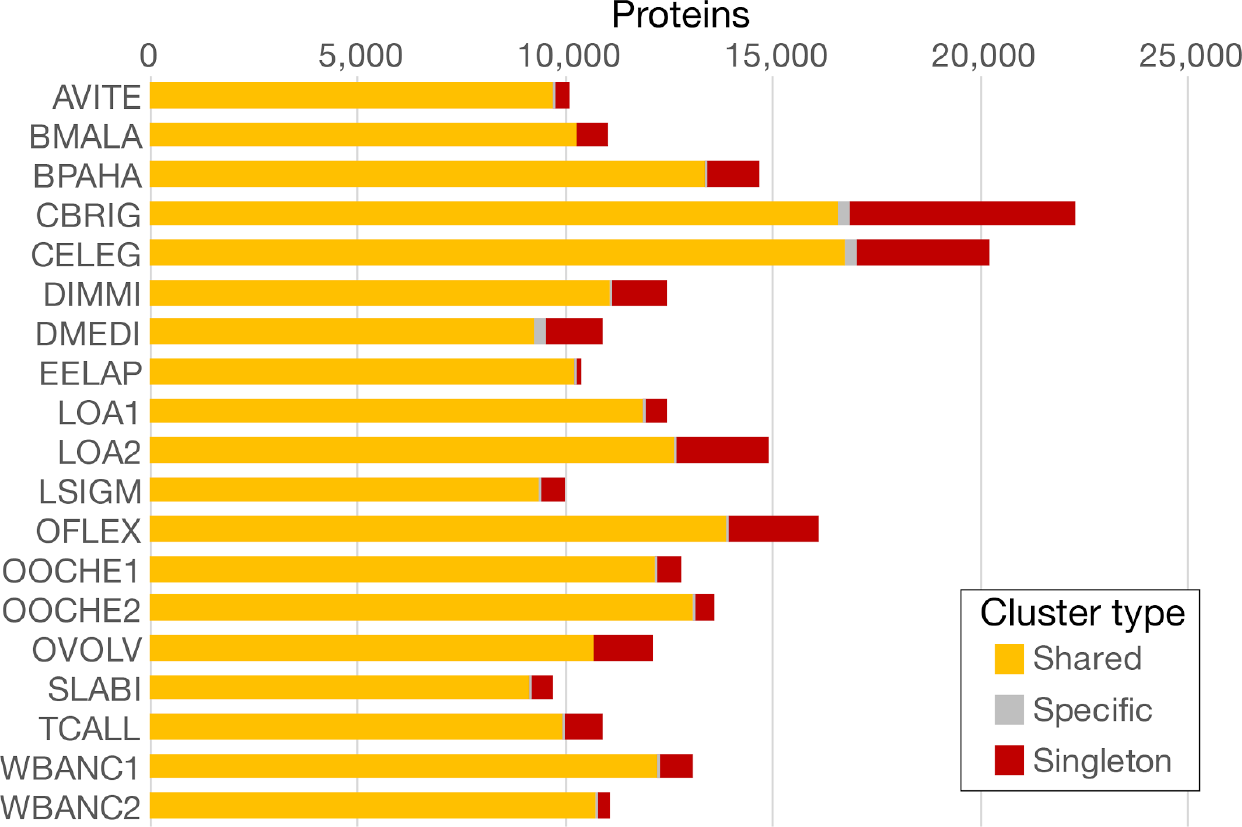
Count of proteins by the type of cluster they were placed in. “Shared”: clusters containing proteins from multiple taxa, “Specific”: clusters containing 2 or more proteins from a single proteome, “Singleton”: clusters containing a single protein.

### Single-copy orthologue definition

Clusters composed of a single protein from each proteome (*i.e.* putative single-copy orthologues) are useful for downstream phylogenetic analyses (see Figure 3). However, due to the intrinsic difficulties of genome assembly and annotation, the number of single-copy orthologues decreases the more proteomes are included in the clustering. To compensate for this, KinFin can identify 'fuzzy' single-copy orthologue clusters using the parameters `target_count` (target number of copies per proteome, default 1), `target_fraction` (proportion of proteomes at `target_count`), and lower/upper counts for proteomes outside of `target_fraction` (`min` and `max`).

**Figure 3.**
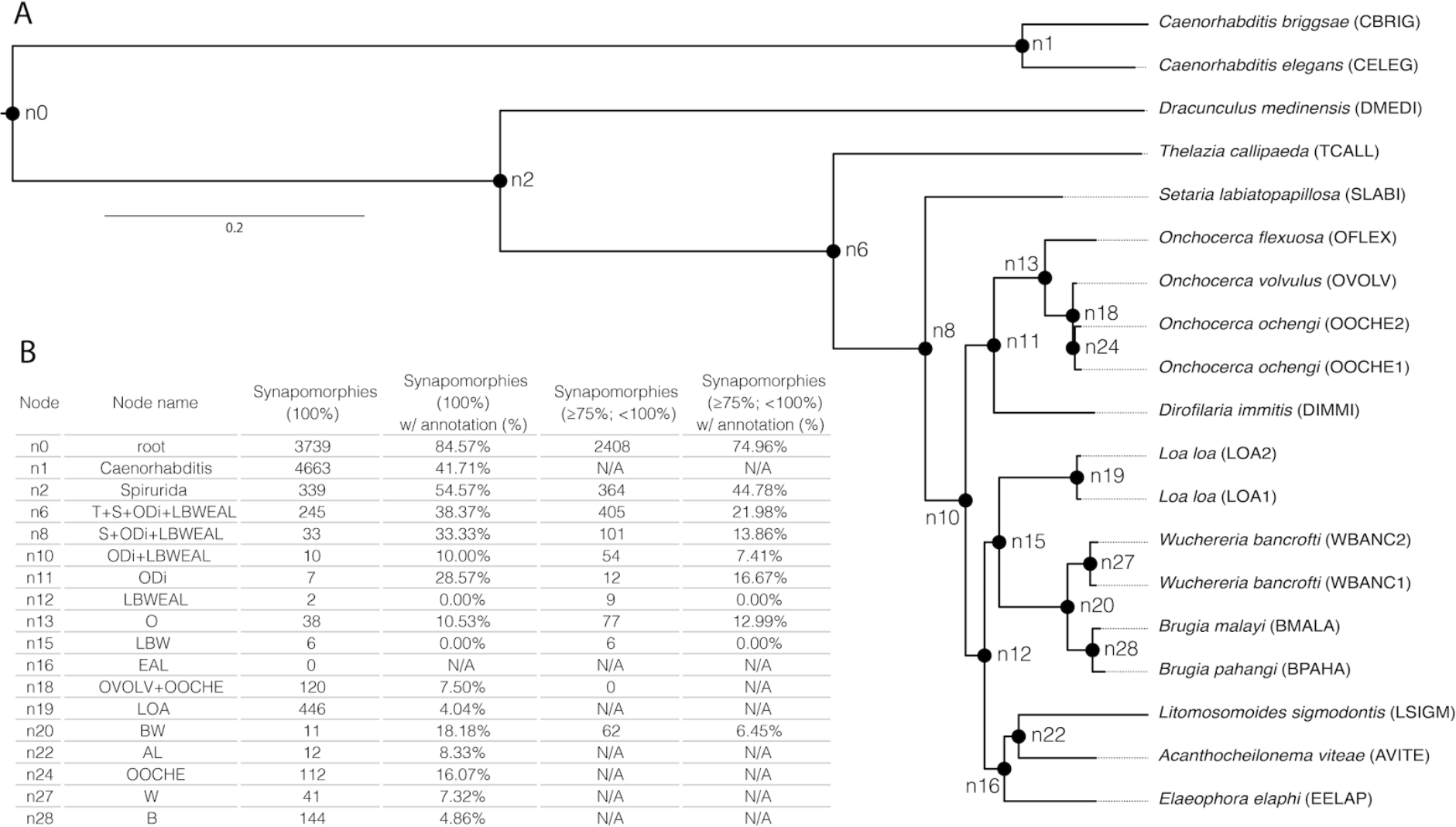
A: Phylogenetic tree based on 781 single-copy orthologues. Non-parametric bootstrap support for all branches is 100. Internal nodes are labelled. B: Table summarising “complete_presence” synapomorphic clusters (100% taxon-coverage) and “partial absence” synapomorphic clusters (75% ≤ taxon-coverage < 100%) and the percentage for which a representative functional annotation could be inferred. “N/A” is used for cases in which nodes are ancestors of less than four taxa or when percentage of functional annotation could not be calculated due to lack of clusters.

### Rarefaction curves

The concept of the pangenome is frequently used in microbial genomics to describe all the genes, both core and accessory, that are found in the varied genomes of a species. The size of the pangenome can be visualised using rarefaction curves, and KinFin deploys this framework to visualise the size of the pan-proteome of the different arbitrary sets defined by the user. Curves are calculated by repeated, random sampling of the proteomes in each arbitrary set and cumulative summation of novel non-singleton clusters (see Figure 4).

**Figure 4.**
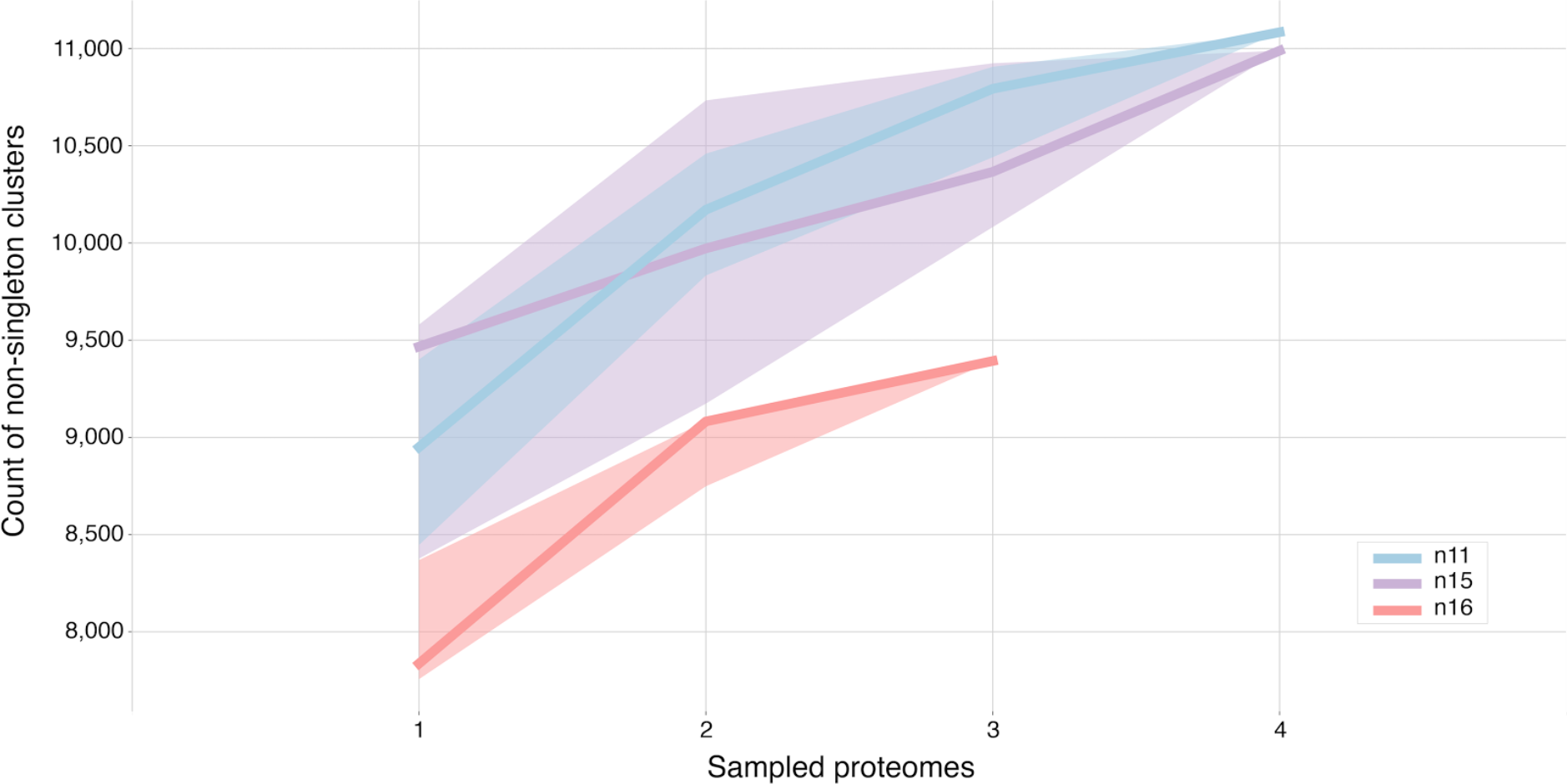
Rarefaction curve of proteomes within sets defined by major clades within the onchocercid nematodes. n 1 1 = node 1 1 (*Dirofilaria immitis* and *Onchocerca* species); n15 = node 15 (*Wuchereria bancrofti, Brugia* species and *Loa loa*); n 1 6 = node 16 (*Litomosoides sigmodontis, Acanthocheilonema viteae* and *Elaeophora elaphi*).

### Pairwise protein count representation tests

For user-defined attributes involving two (or more) taxon sets, pairwise representation tests of protein counts are computed for clusters containing proteins from each taxon set using either two-sided Mann-Whitney U tests (default), Welch’s t-tests, or a Student’s t-tests. From this, clusters “enriched” or “depleted” in count in one set compared to another can be identified. It should be noted that the statistical tests test for non-homogeneity of the distributions of protein counts between the sets and, due to limited “sample size”, might not achieve significance even when counts differ substantially between sets. In addition to text outputs, volcano plots (log2-fold change in means *versus* test p-value) are drawn (see Figure 5). As a visual aid, horizontal lines are drawn at p-values 0.05 and 0.01 and vertical lines at |log2-fc(means)| = 1 and 95%-percentile of log2-fold changes in mean.

**Figure 5.**
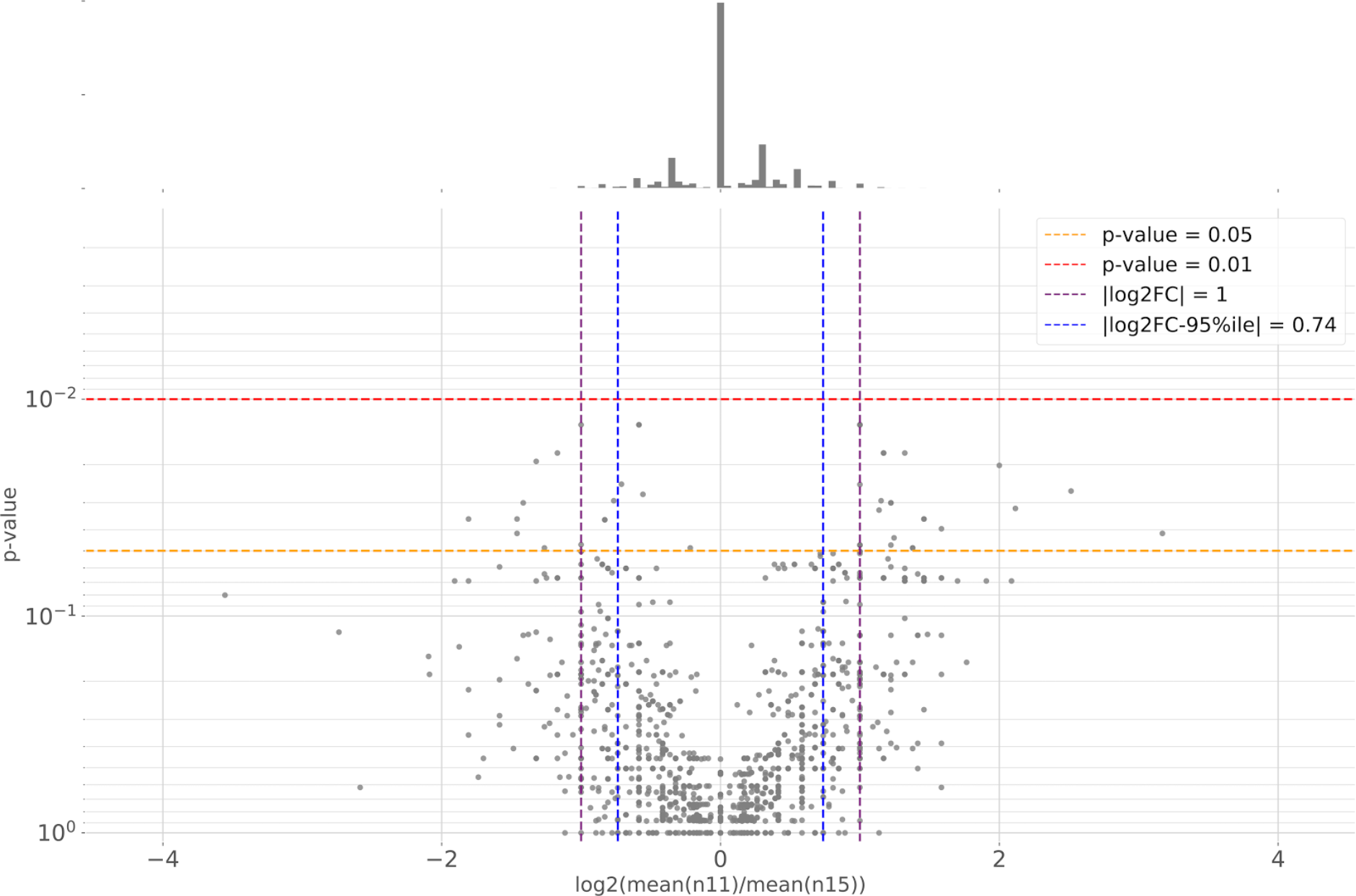
Volcano plot of the protein count representation tests for clusters shared between taxa at node n 1 1 (*Dirofilaria immitis* and *Onchocerca* species) and node n 15 (*Wuchereria bancrofti, Brugia* species and *Loa loa*).

### Analyses based on functional annotation and protein length data

KinFin integrates functional annotation and protein length data into analyses. If the necessary input files are provided, KinFin generates output files tabulating mean and standard deviation of sequence lengths, domain and Gene Ontology (GO) term entropy within clusters, and the fraction of proteins per cluster which are putatively secreted (based on SignalP_Euk annotation). Additionally, for each cluster all matching domains and inferred GO terms are listed with description and information regarding their frequency within both proteins and proteomes in the cluster.

While inference of functional annotation of a protein is relatively straightforward, no clear standards exist for inferring representative functional annotation of clusters of proteins. We provide a script with parameters `domain-taxon coverage` (minimum fraction of taxa in cluster that have at least one protein annotated with that domain) and `domain-protein coverage` (minimum fraction of proteins in cluster annotated with that domain), to grant users fine control over cluster functional annotation.

### Analyses based on phylogeny

Analysis of clusters in a phylogenetic context permits the identification and quantification of clusters that are unique innovations of certain monophyletic groups (*i.e.* synapomorphies). Based on a user-defined tree topology, KinFin identifies synapomorphic clusters at nodes using Dollo parsimony, requiring that only the proteomes under a given node are members of the cluster and that at least one taxon from each child node is a member. Since Dollo parsimony does not penalise multiple losses, KinFin classifies synapomorphies into `complete_presence` and `partial_absence` subgroups. The outputs include lists of synapomorphies and apomorphies ('singleton' and 'non-singleton' proteome-specific clusters) and detailed description of synapomorphic clusters at each node. Prominent or consistent functional annotation can be mapped onto synapomorphic clusters, filtered by `node-proteome coverage` (minimum presence of proteomes as fraction of total proteomes under the node), `domain-proteome coverage` (minimum proportion of proteomes represented by proteins with a specific domain annotation), and domain-protein coverage (minimum proportion of all the proteins annotated with a specific domain).

### Analyses of clusters containing genes of interest

Output of protein clustering analysis often serves as substrate for the identification of homologues of genes of interest from a model species in the target species. KinFin is distributed with a script which takes as input a list of protein IDs or of gene IDs (to obtain all isoforms or only the isoforms included in the clustering) and writes tables indicating the counts of proteins in each cluster and their representative functional annotations.

### Output

KinFin generates output folders for each relevant column in the config file and writes overall metrics for all taxon sets, detailed metrics for each cluster and results of pairwise representation test, draws the rarefaction curve and volcano plots, and lists clusters classified as “true” and “fuzzy” single-copy orthologues. Resulting text files can easily be interrogated using common UNIX command line tools or spreadsheet software.

## KinFin in action: filarial nematodes parasitic in humans and other vertebrates

We illustrate some of the main functionalities of KinFin by addressing questions regarding the biology of filarial nematodes. Filarial nematodes (Onchocercidae) include many species of medical and veterinary interest and the phylogenetic relationships among them remain under debate (Park et al. 2011; Nadler et al. 2007), with the current NCBI reference taxonomy likely to be incorrect. We analysed the proteomes of sixteen species: ten filarial nematodes, three related spirurid nematodes and two *Caenorhabditis* species. *Caenorhabditis* were included because of the quality of available structural and functional annotations. For three species, two independent assemblies and proteome predictions were included.

We will use KinFin to generate a robust multi-locus alignment and phylogeny, and then incorporate this tree into KinFin analyses of synapomorphies and other features of groups of filaria. What gene family and functional differences characterise the different groups? Finally, we will use sets of *Caenorhabditis elegans* genes implicated in pathways of interest to highlight orthology and paralogy in the filarial nematodes.

### KinFin analysis protocol

Here we summarise the protocol used to generate the analyses reported. Detailed protocols including programme versions and command lines are given in Supplementary Methods.

The proteomes detailed in Table 1 were downloaded from WormBase parasite (WBPS8) (Howe, Bolt, Shafie, et al. 2016; Howe, Bolt, Cain, et al. 2016) and https://ngenomes.org. For currently unpublished genomes, manuscripts are in preparation giving details of assembly and annotation strategies. The files were filtered by keeping only the longest isoforms and excluding sequences shorter than 30 residues or containing non-terminal stops. Proteins were functionally annotated using InterProScan (Jones et al. 2014) and clustered using OrthoFinder (Emms and Kelly 2015) at default MCL inflation value of 1.5. using BLASTp (Camacho et al. 2009) similarity data.

**Table 1.**
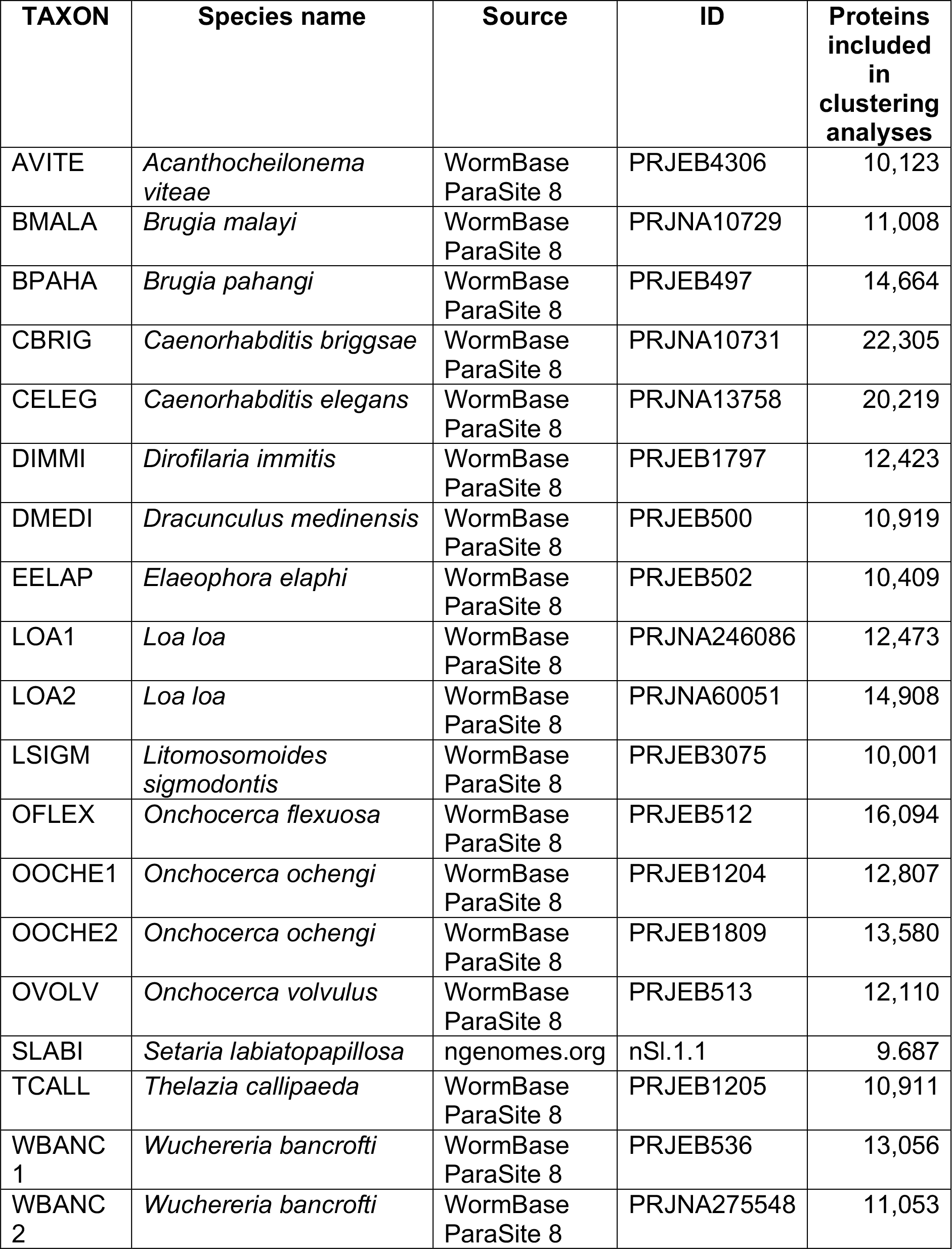
Proteomes used in OrthoFinder clustering and subsequent KinFin analyses.

An initial KinFin analysis identified 781 one-to-one single-copy orthologues. Sequences for these 781 clusters were extracted and aligned using mafft (Katoh and Standley 2013). Alignments were trimmed using trimal (Capella-Gutiérrez et al. 2009), concatenated using FASconCAT (Kück and Meusemann 2010), and analysed using RAxML (Stamatakis 2014) under the PROTGAMMAGTR model of sequence evolution. Non-parametric bootstrap analysis was carried out for 100 replicates.

KinFin was then rerun, providing additional classification in the config file and functional annotation data (Supplementary Dataset 2). The topology of the tree inferred through phylogenetic analysis was provided in Newick format and the two *Caenorhabditis* species were specified as outgroup. The Mann-Whitney-U test was selected for pairwise protein count representation tests. Representative functional annotation was inferred for all clusters, and analysed specifically for subsets of taxa. Genes involved in heme biosynthesis and homeostasis were identified based on representatives from *C. elegans*, and absence of missing genes was confirmed using TBLASTn (Camacho et al. 2009) against the respective genomes. Presence of paralogues was confirmed by manual inspection of gene models on WormBase ParaSite.

## Results

The nineteen proteomes (derived from sixteen species) included 248,750 protein sequences (with a total amino-acid length of 95,162,557 residues) after filtering. OrthoFinder places these into 42,691 clusters, of which 57.97% were singletons (containing 9.95% of protein sequences). The clusters displayed a power-law like frequency distribution, but with a marked deviation from this expectation at a cluster size of 19 (Figure 1). This pattern, although less pronounced, has been observed before for protein databases such as COG (Clusters of Orthologous Groups of proteins) (Unger et al. 2003) and TRIBES (Enright et al. 2003), and has been seen in other datasets analysed with KinFin (Yoshida et al. 2017). These clusters contain a large number of strict (“true”) single-copy orthologues, and many “fuzzy” single-copy orthologues.

KinFin can assist in deciding which of several alternative proteome predictions is more likely to be correct, and identify major differences between alternative predictions. Analysis of between- proteome linkage using Gephi (see Supplementary Results) showed that while some proteome pairs (for example the two from *O. ochengi*) were found to be neighbours, the two *W. bancrofti* proteomes were not placed together, suggesting divergence in prediction. Examination of the distribution of clusters within species can highlight outlier datasets. Both *C. elegans* and *C. briggsae* have higher total protein counts than any of the filarial species and display the highest proportion of singletons (15.7% and 24.4%) (Figure 2). For those species for which two assemblies were analysed, variation in proportion of singletons is most severe for *L. loa* (15.1% vs 4.6%).

The 781 single-copy orthology clusters yielded a robustly supported phylogeny (Figure 3A). Rooting the tree with *Caenorhabditis*, the three non-filarial taxa are recovered in expected positions, with *Setaria labiatopapillosa* most closely related to the onchocercids, followed by *Thelazia callipedia* and *Dracunculus medinensis*. The relationships between the onchocercid taxa is not congruent with the reference NCBI taxonomy, but with a previous analysis using a smaller number of loci (Lefoulon et al. 2016). *Dirofilaria immitis* is recovered as sister to *Onchocerca* species (the clade defined by node n11 in Figure 3A), and *W. bancrofti, Brugia* species and *Loa loa* (node n15) form a clade distinct from *Litomosoides sigmodontis, Acanthocheilonema viteae* and *Elaeophora elaphi* (node n16). We identified 3,887 “fuzzy” single-copy orthologues. These are useful for analyses of proteomes from more distantly related species, where stochastic absence and duplication can severely limit the number of single-copy loci recovered for phylogenetic analyses. Fuzzy orthologues can be used in combination with tools such as PhyloTreePruner (Kocot et al. 2013) which filters out-paralogues and selects appropriate in-paralogues.

We explored the proteomic diversity represented by the three clades within Onchocercidae (Figure 3A, at nodes n11, n15, n16). We defined taxon sets under each of the nodes and used KinFin to generate rarefaction curves for each set (Figure 4). Curves for node n11 (*Dirofilaria immitis* and *Onchocerca* species) and node n15 (W. *bancrofti, Brugia* species and *Loa loa*) show comparable slopes and the number of non-singleton clusters recovered in both is very similar (11,084 clusters for n11 and 10,989 for n15). Fewer unique protein clusters (9,393) were recovered when sampling node n16 (*Litomosoides sigmodontis, Acanthocheilonema viteae* and *Elaeophora elaphi)*. The fact that none of the curves reaches a plateau suggests that their protein space has not been sampled exhaustively.

Of the proteins used in the analysis, 157,873 (63.47%) were annotated with InterPro (IPR) domains. By inferring representative IPR annotations we functionally annotated 12,026 (28.17%) of the clusters. Using the phylogeny inferred from single-copy orthologues (Figure 3 A) we inferred synapomorphies at each node and investigated their representative functional annotations (Figure 3 B). While many clusters are inferred to be synapomorphic for deeper nodes, Onchocercidae and the three groups within Onchocercidae have few synapomorphic gene family births (10 at node n10, 7 at node n11, 6 at node n15 and none at node n16). Of those, only two clusters at node n11 received representative functional annotation: a Chromadorea ALT cluster (OG0007060) and a SOCS box domain cluster (OG0009843). The Chromadorea ALT domain is found across Nematoda and is found in several clusters. *B. malayi* ALT-1, the first described Chromadorea ALT protein (contained in cluster OG0000082), has been proposed as a candidate vaccine target for human lymphatic filariasis (Gregory et al. 2000). The synapomorphic Chromadorea ALT cluster is specific to *Onchocerca* spp. and *D. immitis* and might harbour the same potential for onchocerciasis. SOCS box domains were first identified in proteins involved in suppression of cytokine signalling, and are key regulators of both innate and adaptive immunity (Alexander 2002). Proteins in OG0009843 do not contain any of the other domains usually associated with SOCS, such as SH2 (a combination found in OG0000874 and OG0007539) or Ankyrin repeat-containing domains (a combination found in OG0001559 and OG0015826). However, they may still play an immunomodulatory role during infection as has been suggested for SOCS box proteins in *L. sigmodontis (Godel et al. 2012*). An additional SOCS box cluster is synapomorphic for *Onchocerca* spp.

Definition of taxon sets based on host species (“human” vs “other” vs “outgroup”) recovered 628 clusters specific to filarial nematodes, but none had proteins of more than four out of seven taxa. Of the eight clusters containing proteins of four taxa, six contained only *B. malayi, L. Loa* and *W. bancrofti*. Hence, we found no evidence of systematic convergent adaptation to human hosts in the analysed proteomes of filarial nematodes.

KinFin permits rapid assessment of differences in copy number between species and taxon sets using protein count representation tests. Analysis of clusters shared between taxa either side of the basal split in Onchocercinae (node n11: *Dirofilaria* plus *Onchocerca*, and node n15: other filaria) (Figure 5) identified 10 clusters with extreme differences (see Table S1). Among these was cluster OG0000051, which includes prolyl 4-hydroxylase orthologues, including *Bm*-PHY-1 and *Bm*-PHY-2 which are essential for development and cuticle formation, and have been suggested as potential targets for parasite control (Winter et al. 2013). While all n15-taxa have exactly two paralogues (WBANC2 had only *Wb*-PHY-1, but *Wb*-PHY-2 was located through a TBLASTN search and was present in WBANC1), counts in n11-taxa ranged from 5 (OFLEX) and 14 (OVOLV). We identified 3 additional, singleton prolyl 4-hydroxylase clusters, all from n15 taxa. The number of paralogous prolyl 4-hydroxylases in *D. immitis* and *Onchocerca* species could have negative implications in control measures against this locus.

To demonstrate the utility of KinFin for directed analyses of genes and gene families, we revisited the biology of heme synthesis and transport in the Onchocercidae. This pathway is a target of active investigation for drug development. While most organism can synthesize heme, a complete heme biosynthetic pathway is lacking in all nematodes studied to date (Rao et al. 2005), and proteins of only two of 12 catabolic steps (COX-10 and COX-15) have been described in *C. elegans*. In *C. elegans* multiple heme responsive genes (HRGs) have been characterised (Rajagopal et al. 2008; Chen et al. 2011; Sinclair and Hamza 2015) and orthologues have been identified in *B. malayi* (*Bm*-HRG-1 and *Bm*-HRG-2) and *D. immitis* (Luck et al. 2016). In *C. elegans*, HRG are involved in heme trafficking within the epidermis (HRG-2), to oocytes (HRG-3) and within the intestine (HRG-1/4/5/6). Other ABC transporters in *C. elegans* have been implicated in heme homeostasis (MRP-5, F22B5.4, and ABTM-1) (Severance et al. 2010; González-Cabo et al. 2011; Antonicka et al. 2003). An orthologue of MRP-5 has been described in *B. malayi* (Luck et al. 2016). Several animal parasitic nematodes (including *B. malayi, D. immitis*, and *O. volvulus*) have been shown to harbour a functional ferrochelatase (FECH) acquired through horizontal gene transfer from an alphaproteobacterium (Elsworth et al. 2011; Nagayasu et al. 2013; Wu et al. 2013). Other nematodes have distinct ferrochelatase-like (FECL) homologues which lack the active site. We catalogued homologues of these proteins in our clustering analysis (Figure 6).

**Figure 6.**
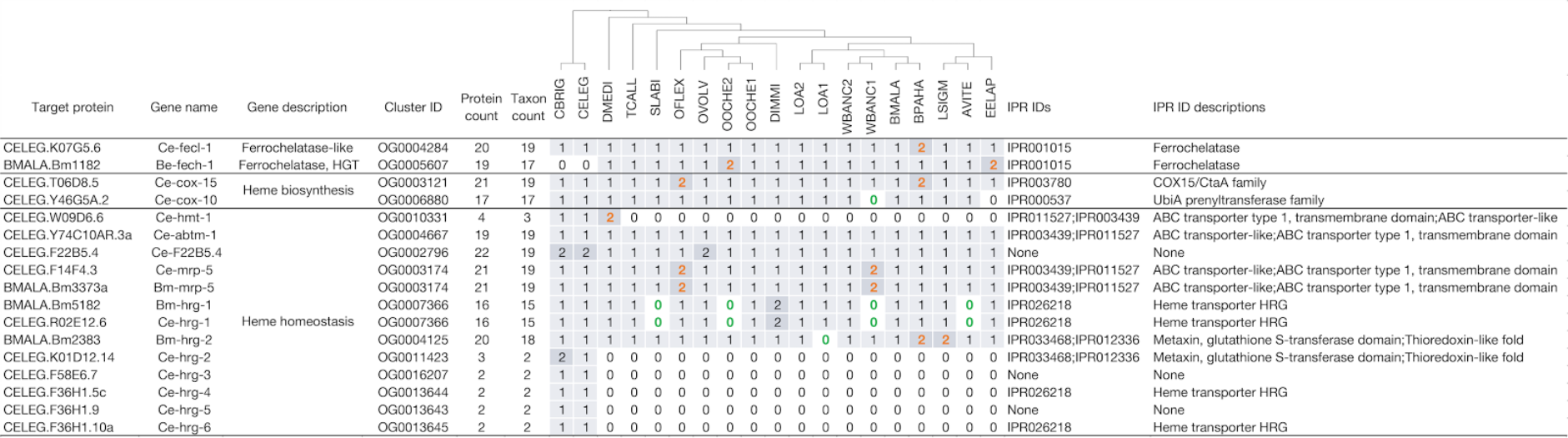
Heme biosynthesis and transport genes. The columns showing counts of proteins per taxon are ordered based on phylogeny (cartoon above). Counts are coloured orange if they result from fragmented predictions and green if the gene was only identified by directed TBLASTN search.

FECL proteins were identified in all species. *B. pahangi* has two FECL proteins while all other taxa have one, but both are located at scaffold borders and may be the result of an assembly artefact. The horizontally-acquired FECH is absent from the *Caenorhabditis* proteomes (and genomes) but present in all the other taxa analysed. Paralogues in one of the *O. ochengi* proteomes are suggestive of misprediction. COX-10 and COX-15 are present in most taxa in the analysis; paralogues in *B. pahangi* and *O. flexuosa* are a result of fragmented assemblies. COX-10 is present in *W. bancrofti* WBANC1 (on scaffold WBA_contig0009713), but the gene was not predicted. COX-10 was not found in *E. elaphi*, which suggests that either the corresponding genomic region was not assembled or that the gene has been lost.

Presence/absence of proteins involved in heme homeostasis show a more complex pattern. *Ce*-HMT-1, an ATP-dependent phytochelatin transporter, is restricted to *Caenorhabditis* spp. and *D. medinensis*. The other ABC-transporter-like proteins (ABTM-1, MRP-5, and F22B5.4) are present across all taxa. For F22B5.4, genuine paralogues are found in both *Caenorhabditis* species and *O. volvulus*. *Ce*-MRP-5 and *Bm*-MRP-5 are located within the same cluster, and apparent paralogues in *O. flexuosa* and *W. bancrofti* WBANC1 derive from predictions located at the ends of scaffolds. While no orthologues of *Ce*-HRG-2/3/4/5/6 were identified, cluster containing *Ce*-HRG-1 included representatives from most species. Missing orthologues of HRG-1 were identified using TBLASTN searches in *Setaria labiatopapillosa* (nSl.1.1.scaf00038), *O. ochengi* OOCHE2 (nOo.2.0.Scaf03259), *W. bancrofti* WBANC1(WBA_contig0001821), and *A. viteae* (nAv. 1.0.scaf00129). The two HRG-1 paralogues in *D. immitis* were identified previously (Luck et al. 2014; Luck et al. 2016). Interestingly, “*Bm*-HRG-2” (Bm2383) is not orthologous to *Ce*-HRG-2 but rather to *Ce*-C25H3.7, an orthologue of human FAXC (failed axon connection).

In summary, all non-*Caenorhabditis* nematodes analysed have a functional FECH orthologous to the one acquired through HGT in *B. malayi*. Proteins responsible for the only two steps in heme biosynthesis described in *C. elegans* are also found in all taxa, apart from COX-10 in *E. elaphi.* The heme transporters HRG-2/3/4/5/6 are (in this analysis) restricted to *Caenorhabditis*, but all the spirurid nematodes analysed have retained HRG-1, a FAXC-like cluster orthologous to *Bm*-HRG-2, and MRP-5, and these may mediate heme transport from the intestine.

## Conclusion

As ever more genomes are sequenced, and our understanding of the diversity of protein space increases, it concomitantly becomes more difficult to see the patterns in complex orthology data. To ease this bottleneck, we have presented KinFin, a tool that takes the output of standard orthology inference toolkits and provides a user-friendly but rich analytical toolkit to review and interrogate orthology clustering. By permitting user definition of custom taxon sets, KinFin can be used to highlight changes in presence or membership of orthologue groups associated with either taxonomy or phenotypes of interest. Its reliance on standard input file formats and explicit parameters makes integration in comparative genomics projects easy, and thus promotes transparent and reproducible analysis of clustered protein data. KinFin is under regular maintenance and we welcome user feedback. Further development will involve a HTML user interface to access and interact with the generated output files.

We presented some of the main capabilities of KinFin through the analysis of proteomes of filarial nematodes and outgroup species. By extracting single-copy orthologues we resolved the phylogenetic relationships between filarial nematodes. We explored synapomorphic clusters and their functional annotations across the phylogeny and identified putative gene families of interest. Through definition of (phylogenetic) taxon sets we explored and visualised proteomic diversity across key clades, and analysed differences in protein counts between the sets. To illustrate targeted analysis of homologues of genes of interest, we analysed clusters containing proteins involved in heme metabolism and homeostasis using characterised orthologues from model organisms.

One of the advantages of KinFin is the speed of computation. Analyses reported here took 40 seconds (basic KinFin analysis) and 4 minutes (advanced KinFin analysis) on a MacBookPro (15-inch, 2016; 2.9Ghz I7, 16Gb RAM). While addition of taxa to the clustering and taxon sets to the config file will increase computation time, we think run time is within acceptable bounds for this type of analysis. KinFin’s speed promotes hypothesis exploration, such as comparing alternative phylogenetic topologies, an example of this has been published recently (Yoshida et al. 2017), or contrasting taxon sets.

## Availability and requirements

KinFin is available under GPL3 license from GitHub (https://github.com/DRL/kinfin). Instructions concerning installation and analysis can be found at https://kinfin.readme.io/docs. Supplementary material from this study is deposited on GitHub (https://github.com/DRL/kinfin_manuscript).

## Acknowledgements

We thank members of the Blaxter Nematode and neglected genomics lab in Edinburgh for support, criticism and suggestions. We thank colleagues who supplied materials for nematode genome sequencing, and the Berriman Lab of the Wellcome Trust Sanger Institute and the staff of Wormbase ParaSite for access to genomes. We thank Duncan Berger for testing the software. DRL is supported by a James Hutton Institute/Edinburgh University School of Biological Sciences fellowship.

